# Intracellular copper storage and delivery in a bacterium

**DOI:** 10.1101/2022.01.05.475036

**Authors:** Jaeick Lee, Christopher Dennison

## Abstract

A family of cytosolic copper (Cu) storage proteins (the Csps) are widespread in bacteria. The Csps can bind large quantities of Cu(I) via their Cys-lined four-helix bundles, and the majority are cytosolic (Csp3s). This is inconsistent with the current dogma that bacteria, unlike eukaryotes, have evolved not to maintain intracellular pools of Cu due to its potential toxicity. Sporulation in *Bacillus subtilis* has been used to investigate if a Csp3 can store Cu(I) in the cytosol for a target enzyme. The activity of the Cu-requiring endospore multi-Cu oxidase *Bs*CotA (a laccase) increases under Cu-replete conditions in wild type *B. subtilis*, but not in the strain lacking *Bs*Csp3. Cuprous ions readily transfer from *Bs*Csp3, but not from the cytosolic copper metallochaperone *Bs*CopZ, to *Bs*CotA *in vitro* producing active enzyme. Both *Bs*Csp3 and *Bs*CotA are upregulated during late sporulation. The hypothesis we propose is that *Bs*Csp3 acquires and stores Cu(I) in the cytosol for *Bs*CotA.

## Introduction

Copper (Cu) is essential for most organisms, but use of this metal ion is associated with significant risks due to its potential toxicity. The availability of Cu is restricted by the presence of high-affinity sites in both eukaryotes (1) and prokaryotes (2). Import, cytosolic handling, trafficking to different locations, and storage have all been characterised in eukaryotic cells (3). In bacteria, some of these processes are either not thought to occur, or are not yet fully understood. For example, the plasma membrane protein CcoG, which reduces Cu(II) to the preferred intracellular oxidation state (Cu(I)) has only recently been identified in bacteria as a cytochrome oxidase (COX) assembly factor (4). The reduction of Cu(II) prior to import into eukaryotic cells has been known to occur for many years (3, 5). Excess Cu(I) is removed from the cytosol by probably the best-studied component of bacterial Cu homeostasis; a Cu-transporting P-type ATPase (CopA), which can be assisted by the cytosolic Cu metallochaperone CopZ (Figure 1a) (3, 6-9). Toxicity has been shown to involve Cu(I) binding in place of the native metal in cytosolic iron-sulfur (Fe-S) cluster-containing proteins (10), and Cu catalyses ROS formation (3, 7, 8). The intracellular damage that Cu can cause, and the current dearth of intracellular Cu-requiring enzymes (11), has resulted in a prevailing view that bacteria have evolved not to use Cu in the cytosol (8, 11). However, there is no *a priori* reason why bacteria, like eukaryotes, cannot utilise Cu in this compartment if mechanisms are available to enable its safe handling, i.e. by ensuring tight chelation and specific delivery. The presence of cytosolic Cu storage proteins (Csps) that can bind large quantities of Cu(I) with high affinity (12-15), provides a possible route for intracellular Cu use in bacteria.

**Figure 1.**
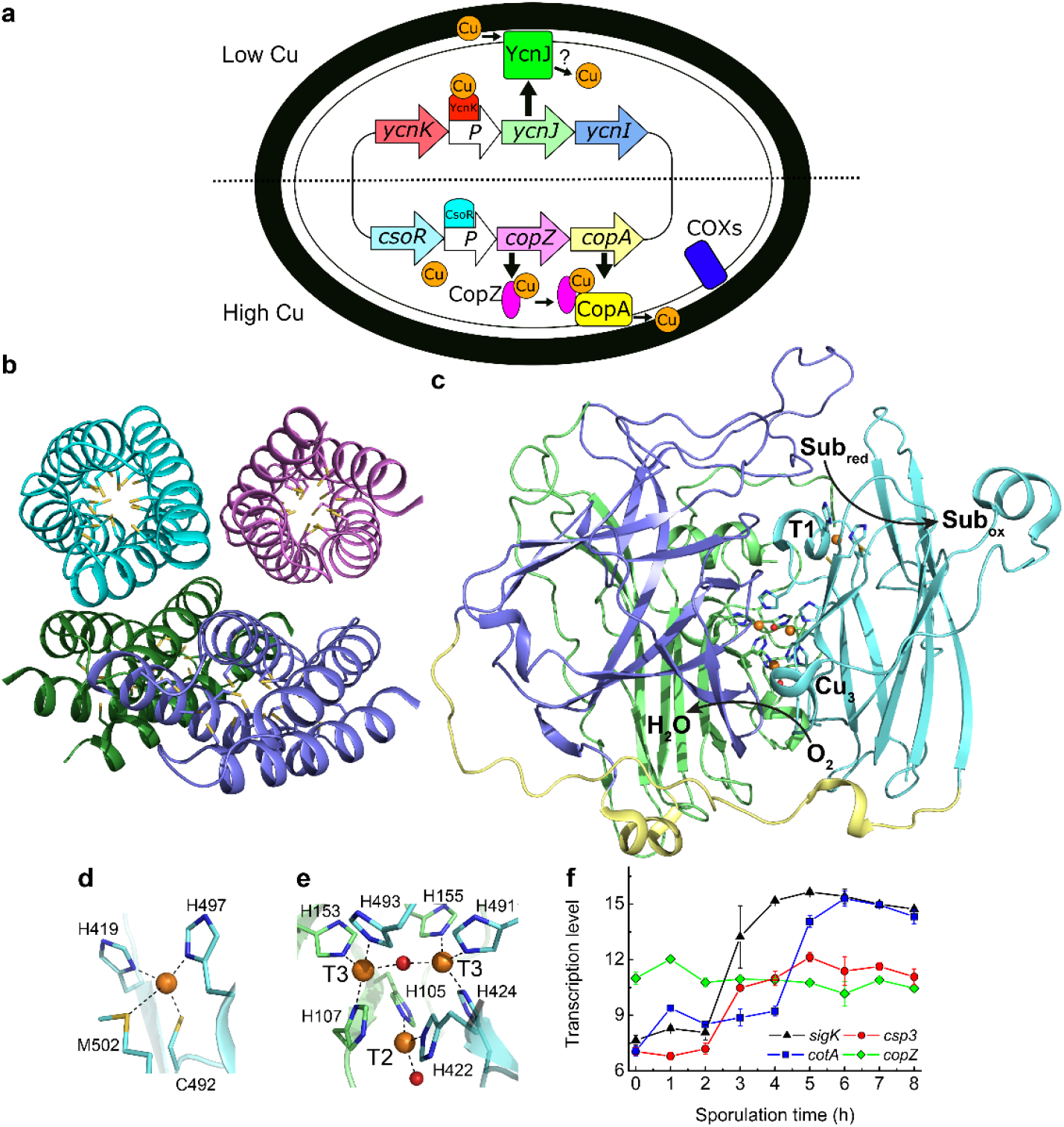
Copper handling, a cytosolic Cu(I) storage protein, a Cu-requiring enzyme, and their transcription during sporulation in *B. subtilis*. (a) An overview of Cu homeostasis in *B. subtilis* including Cu (orange circles, oxidation state undefined) export by CopA and CopZ (regulated by CsoR) (19), and import by YcnJ (regulated by YcnK) (20, 22). YcnI is also membrane bound and binds Cu(II) *in vitro*, but its role in Cu homeostasis is unclear (23). The only currently known Cu-requiring enzymes in vegetative *B. subtilis* cells are two cytochrome oxidases (COXs) located on the plasma membrane (11, 24). (b) The crystal structure of Cu(I)-free *Bs*Csp3 (PDB: 5FIG), a tetramer of four-helix bundles each with 19 Cys residues pointing into their cores enabling the binding of up to ∼20 Cu(I) ions per monomer (13). (c) The crystal structure of the endospore multi-Cu oxidase (a laccase) *Bs*CotA (PDB: 1W6L, (27)) with domains 1, 2 and 3 coloured green, slate and cyan, respectively (the linking regions are yellow). Substrates are oxidized (Sub_red_ to Sub_ox_) at the T1 Cu centre with electrons passed to the T2/T3 trinuclear (Cu_3_) cluster where oxygen is reduced to water. Detailed views of the T1 Cu site (d) and the Cu_3_ cluster (e), with the side chains of coordinating residues represented as sticks, Cu ions as orange spheres and the oxygen atoms of water (bound to the T2 Cu) and hydroxide (bridging the T3 Cu ions) ligands as red spheres in (c-e). (f) Transcription profiles (29) of the *sigK* (σ^K^, which facilitates spore coat protein expression, black triangles), *csp3* (red circles), *cotA* (blue squares) and *copZ* (green diamonds) genes during sporulation.

The Csps were first identified in Gram-negative bacteria that oxidize methane (12). These methanotrophs can possess different Csp homologues, all having many Cys residues lining the cores of their four-helix bundles, enabling the binding of large quantities of Cu(I) ions (12-14). A Csp exported from the cytosol (Csp1) can store up to 52 Cu(I) ions per tetramer for the particulate (membrane-bound) methane monooxygenase (pMMO) in the model methanotroph *Methylosinus trichosporium* OB3b (*Mt*Csp1) (12). *Mt*Csp1 is upregulated (16) at the Cu concentrations required for methane oxidation by pMMO in switchover methanotrophs, which can use a soluble Fe MMO when Cu is limiting (17). However, a cytosolic Csp homologue (*Mt*Csp3) is not upregulated with pMMO in *M. trichosporium* OB3b (16).

The Gram-positive bacterium *Bacillus subtilis* is an excellent model system for investigating the role of a Csp3, as its Cu homeostasis system is well characterised (Figure 1a) (7, 18-23). The *copZA* operon (Cu efflux machinery, *vide supra*) and its Cu-sensing repressor CsoR (7, 18, 19, 21) are probably the best-studied components. The membrane protein YcnJ is upregulated under Cu-limiting conditions, controlled by the suggested repressor YcnK (20, 22), and has been proposed to play a role in Cu acquisition (Figure 1a). The membrane-anchored YcnI is part of the same (*ycnKJI*) operon and is also regulated by YcnK (22). The soluble domain of YcnI binds Cu(II) *in vitro*, and this protein has been suggested to function as a Cu metallochaperone (23). Cytosolic Cu(I) could be safely stored in *B. subtilis* by the Csp3 homologue (*Bs*Csp3) whose core is lined with 19 Cys residues (Figure 1b), which enable the binding of ∼80 Cu(I) ions per tetramer *in vitro* (13). Overexpressing *Bs*Csp3 in the cytosol of a Gram-negative heterologous host (*Escherichia coli*) allows growth at otherwise harmful Cu concentrations, and the protein can acquire Cu(I) in the presence of both CopA and CopZ (15).

Only two families of Cu enzymes are currently known to be present in *B. subtilis*; COXs located on the plasma membrane and the multi-Cu oxidase (MCO; a laccase) *Bs*CotA (11, 24-27). The latter is an outer spore-coat (endospore) enzyme (28) that possesses the typical type 1 (T1), 2 (T2) and 3 (T3) Cu sites of an MCO (27), which are involved in the catalytic cycle (see Figure 1c-e). It produces a melanin-like pigment thought to provide spores with protection against hydrogen peroxide and UV light (25, 28). *Bs*CotA is upregulated during the latter stages of sporulation, as is *Bs*Csp3 (Figure 1f) (29).

*Bs*Csp3 does not provide resistance to Cu toxicity. We have therefore tested the hypothesis that *Bs*Csp3 safely stores Cu(I) ions in the cytosol for a Cu-requiring enzyme by investigating the effect of gene deletion on the activity of *Bs*CotA in spores grown under Cu limiting and replete conditions. The data obtained indicate a direct role for *Bs*Csp3 in ensuring the maximum activity of *Bs*CotA. The ability of *Bs*Csp3 to activate *Bs*CotA has been confirmed by the *in vitro* transfer of Cu(I) between these proteins. A model for how *Bs*CotA acquires Cu(I) from *Bs*Csp3 during sporulation is proposed. This is the first example showing that an enzyme can acquire Cu in the cytosol of a bacterium as well as identifying the partner protein.

## Results

### Is *Bs*Csp3 involved in combating Cu toxicity in *B. subtilis*?

The presence of a protein with a high capacity for Cu(I) in the cytosol of *B. subtilis* (12-14) would suggest a role in helping to prevent the problems caused by excess Cu (10, 15). The toxicity of Cu to bacteria is highlighted by the influence increasing Cu concentrations in media has on the growth of wild type (WT) *B. subtilis* (Supplementary Figure 1). At higher Cu levels cells grow extremely slowly, with a very small increase in the absorbance/OD observed only after more than 6 h at 2 mM Cu, and this coincides with elevated intracellular Cu concentrations (Supplementary Figure 2). Very similar growth and Cu accumulation results are obtained for the strain (Δ*csp3*) lacking the *csp3* gene (Supplementary Figures 1 and 2). These data exhibit an overall likeness to work we reported previously (13), particularly at up to 12 h growth. However, the response to specific Cu concentrations varies, indicating the amounts of Cu added to media must differ. In the present work we are sure of the Cu concentrations in media, having carefully quantified all Cu(II) stocks before their addition, and the reported values in the previous study (13) appear to be too high. The growth studies reported herein demonstrate that *Bs*Csp3 is not involved in helping prevent the harmful effects on *B. subtilis* caused by elevated Cu levels. Therefore, the protein does not have a function like eukaryotic Cys-rich metallothioneins (3).

### Using sporulation to determine the function of *Bs*Csp3

The *csp3* and *cotA* genes are both upregulated (29) at similar stages during sporulation (Figure 1f), and *Bs*CotA is the only known Cu-requiring enzyme present in spores. We have therefore studied whether *Bs*Csp3 stores Cu(I) for *Bs*CotA. This enzyme binds four Cu ions (Figure 1c-e) and oxidises the laccase substrate 2,2′-azino-bis(3-ethylbenzothiazoline-6-sulfonic acid) (ABTS) *in vitro* and in spores (26). For WT *B. subtilis* spores the ability to oxidise ABTS increases approximately two-fold when 50 μM Cu is added to media during sporulation (Figure 2a and Supplementary Figure 3a,b). Incubation with Cu(II) only enhances the activity of purified WT spores obtained in the absence of added Cu (Supplementary Figure 4 and Table 1), consistent with previously reported data (26). This indicates that unless supplemented, sporulation media does not contain sufficient Cu (∼0.4 μM) to fully metallate all of the *Bs*CotA produced. However, the addition of 50 μM Cu, a non-toxic concentration, during growth enables *B. subtilis* to obtain enough of the metal to generate only fully Cu-loaded *Bs*CotA in the endospore.

**Figure 2.**
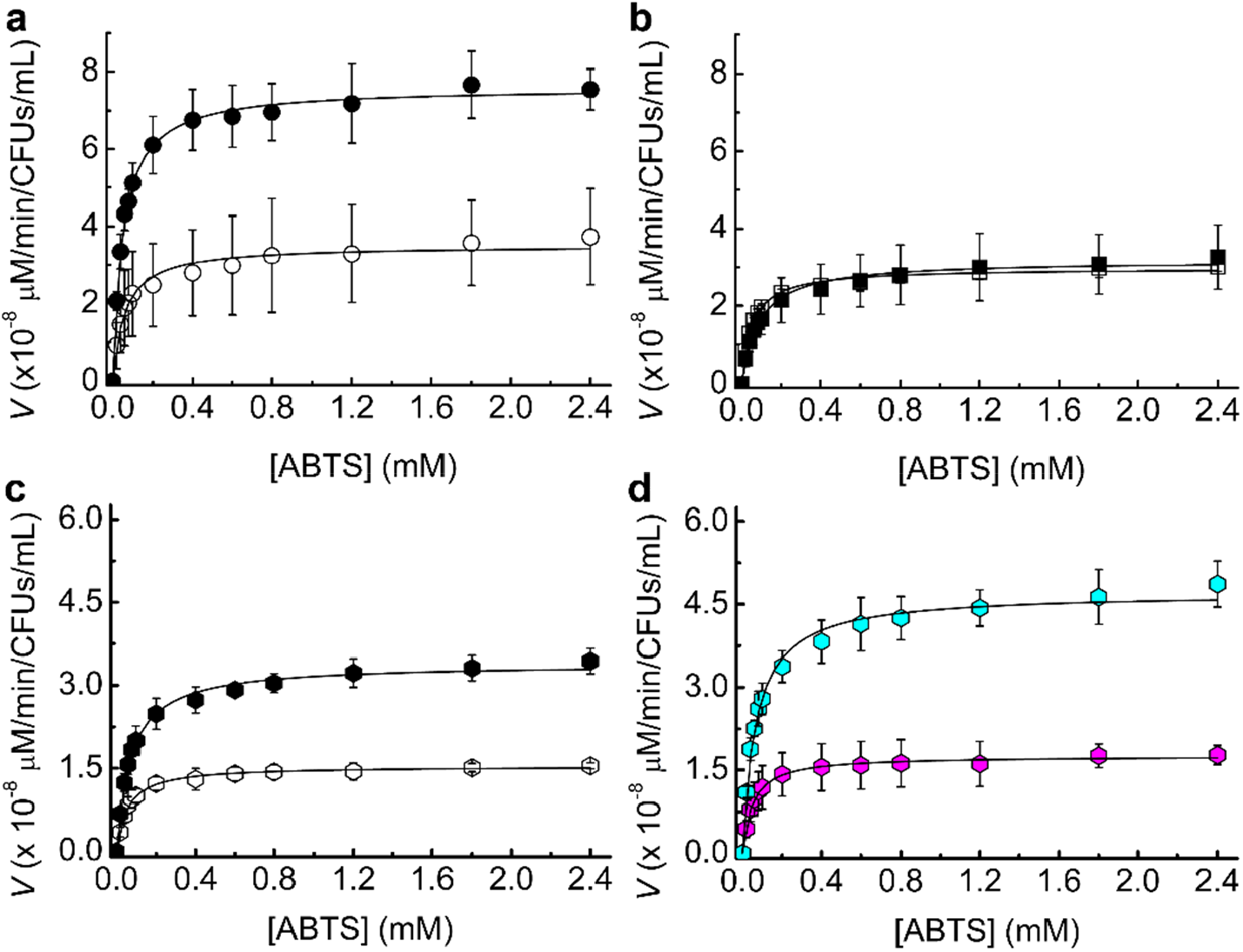
The influence of Cu levels and deleting the *csp3* gene on *Bs*CotA activity in *B. subtilis*. Michaelis-Menten plots of *Bs*CotA activity for heated purified spores from WT (a), Δ*csp3* (b), and the complemented Δ*csp3* (c, d) strains. Spores (a, b and c) were produced in Difco sporulation media plus no (open symbols) and 50 μM (black filled symbols) added Cu(NO_3_)_2_. (d) For the complemented Δ*csp3* strain sporulation was also carried out in the presence of 1 mM IPTG with either no (magenta) or 50 μM (cyan) added Cu(NO_3_)_2_. The plots from which the initial rates for WT and Δ*csp3 B. subtilis* spores were obtained are shown in Supplementary Figure 3. Averages and standard deviations from kinetic measurements in 100 mM citrate-phosphate buffer pH 4.0 using three different sets of spores are shown.

The *Bs*CotA activity of Δ*csp3 B. subtilis* spores grown without added Cu is similar to that for WT spores produced under the same conditions (Figure 2b and Supplementary Figure 3c,d). However, unlike for WT *B. subtilis*, supplementing media with Cu during sporulation has no effect on *Bs*CotA activity for the Δ*csp3* strain. Similar to the WT data, incubation with Cu only significantly increases the activity of purified Δ*csp3* spores grown in the absence of added Cu (Supplementary Figure 4 and Table 1). These results demonstrate that *Bs*Csp3 plays a role in storing Cu for *Bs*CotA, particularly under Cu-replete conditions. Some *Bs*CotA activity remains for Δ*csp3 B. subtilis* spores, and an alternative mechanism of Cu transfer to *Bs*CotA must exist, which could also be responsible for the activity observed in the WT strain under Cu-limiting conditions.

To confirm that *Bs*Csp3 supplies Cu(I) to *Bs*CotA *in vivo*, the Δ*csp3* strain was complemented by introducing an inducible copy of the *csp3* gene at a different location (the *amy*E locus). The highest *Bs*CotA activity is obtained for spores of this strain grown in the presence of isopropyl β-D-thiogalactopyranoside (IPTG, the inducer) and Cu (Figure 2c,d). Activity is almost three-fold greater than without their addition, similar to the increase caused by Cu in WT *B. subtilis* spores (Figure 2a). Elevated activity is observed for spores from the complemented strain grown with Cu, but not IPTG, present in the media (Figure 2c). To ensure this effect is due to leaky expression, a well-established feature of the promoter used [for example, see ref. 30], a control Δ*csp3* strain was constructed (see Material and Methods). In this case, the ability to oxidise ABTS is hardly influenced by the presence of Cu and IPTG (Supplementary Figure 5). Therefore, the increased *Bs*CotA activity for the complemented Δ*csp3* strain is due to the IPTG-inducible copy of the *csp3* gene.

### Cu(I) transfer from *Bs*Csp3 to *Bs*CotA *in vitro*

The above data support the hypothesis that *Bs*Csp3 can store Cu(I) in the cytosol under Cu-replete conditions, which is used to metallate *Bs*CotA. To further test this idea, the transfer of Cu(I) from *Bs*Csp3 to inactive apo(Cu-free)-*Bs*CotA has been studied *in vitro*. We have previously found (12, 13) that Cu(I) removal from Csp3s by a large excess of high-affinity Cu(I) ligands is slow (∼60% removal in 24 h for *Bs*Csp3 using bathocuproine disulfonate, see Supplementary Figure 6 and Table 2). The transfer of Cu(I) from *Bs*Csp3 to *Bs*CotA occurs quickly when using a 10-fold excess of Cu(I) in the storage protein over sites in the enzyme, and >50% maximum activity is achieved within 6 h (Figure 3 a,b and Figure 4a). A control experiment was performed studying activation of as-isolated *Bs*CotA with Cu(I) in the absence of *Bs*Csp3, which is faster, but still takes 4 h to complete.

**Figure 3.**
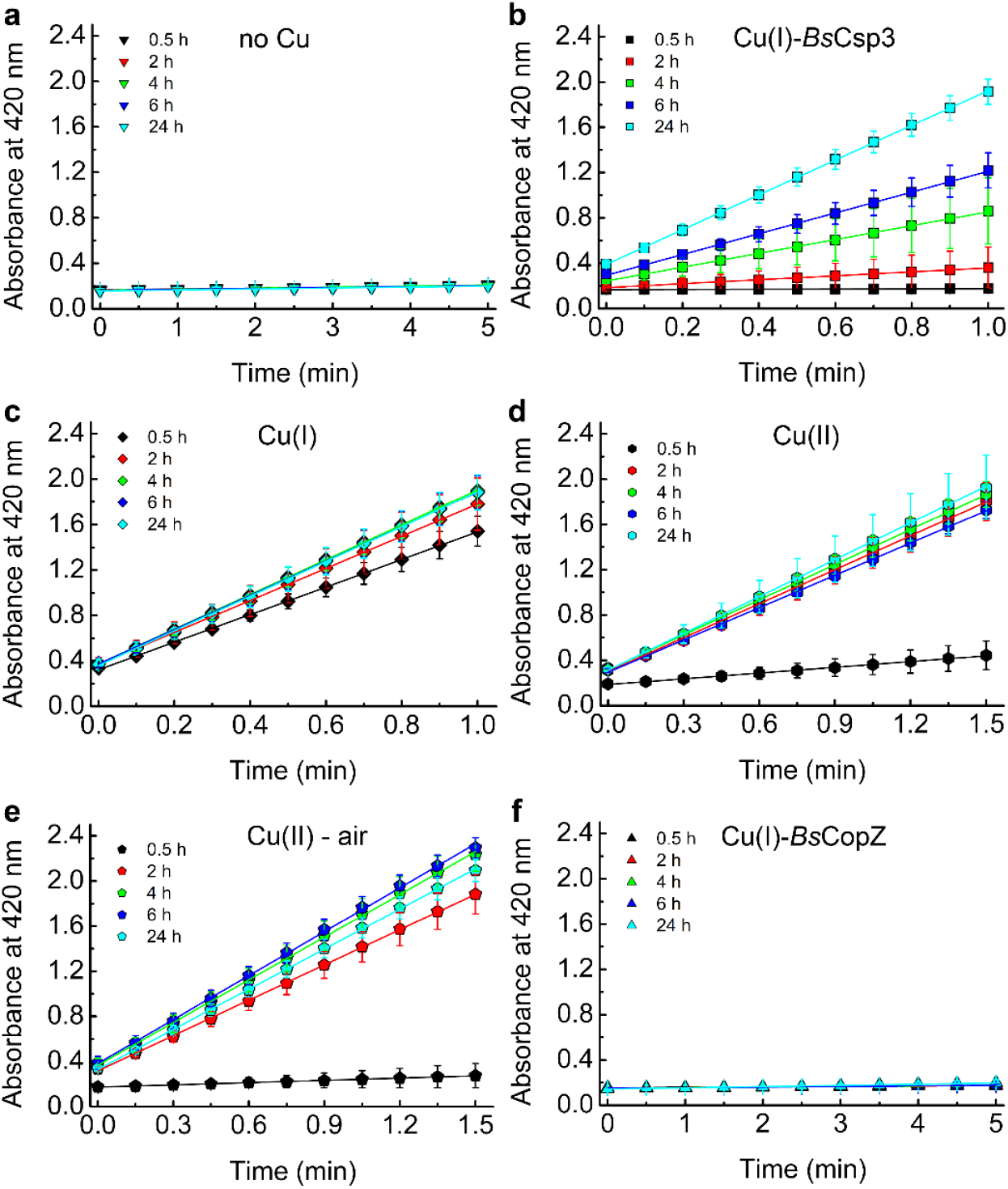
Kinetic analysis of the activity of *Bs*CotA after incubation with Cu from difference sources. Plots of absorbance at 420 nm against time for the reaction with 2.4 mM ABTS of mixtures of apo-*Bs*CotA incubated with buffer (a), Cu(I)-*Bs*Csp3 (b), Cu(I) (c), Cu(II) (d and e) and Cu(I)-*Bs*CopZ (f) for 0.5, 2, 4, 6 and 24 h. Mixtures were incubated in 20 mM HEPES pH 7.5 plus 200 mM NaCl (the buffer used in a), and *Bs*CotA activity was measured 100 mM citrate-phosphate buffer pH 4.0. All data (averages from three independent experiments with error bars showing standard deviations) are from mixtures incubated under anaerobic conditions, apart form in **e** when *Bs*CotA and Cu(II) were incubated in air. The resulting activity data at these and other incubation times are shown in Figure 4.

**Figure 4.**
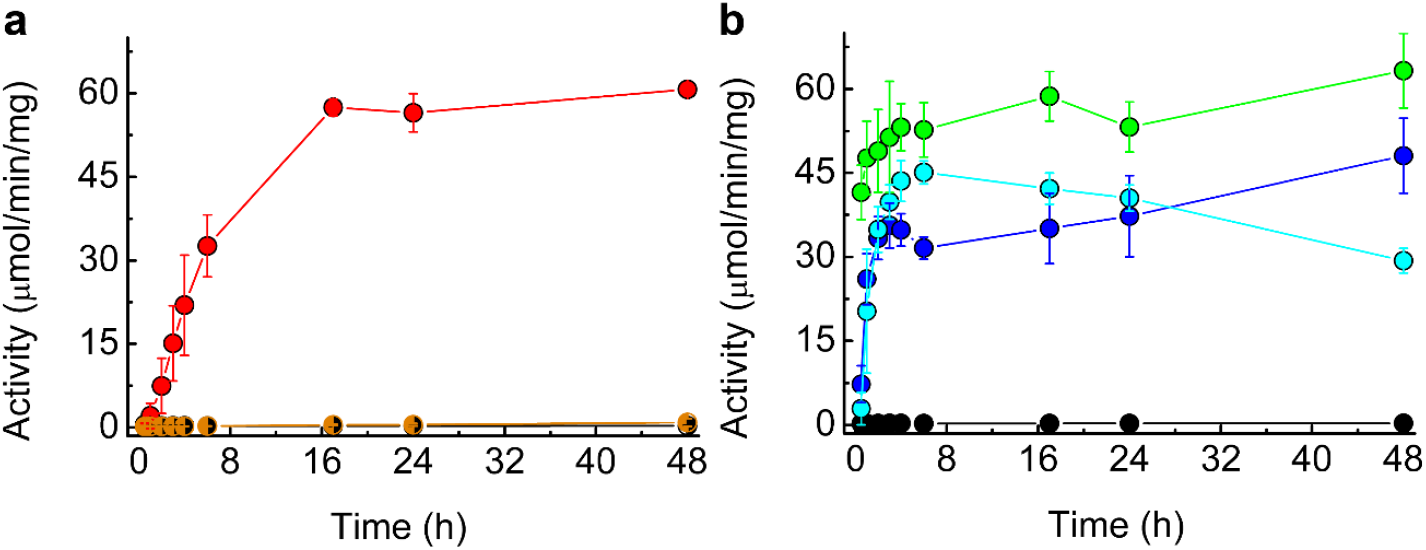
Cu(I) transfer from *Bs*Csp3 to *Bs*CotA. (a) Plots of activity (average from three independent experiments with error bars showing standard deviations) against incubation time of as-isolated *Bs*CotA incubated with Cu(I)-*Bs*Csp3 (red circles), Cu(I)-*Bs*CopZ (half-orange circles) and buffer alone (half-black circles) for up to 48 h under anaerobic conditions. The activity data for as-isolated *Bs*CotA plus buffer alone (black circles) are also shown in (b), as well as the results from control experiments in which the enzyme was incubated with Cu(I) (green circles) and Cu(II) under anaerobic conditions (blue circles), and also with Cu(II) in air (cyan circles). The kinetic data at selected time points are shown in Figure 3.

A further control was carried out using Cu(II) and metalation of the enzyme is slower than for Cu(I) both under anaerobic and aerobic conditions (Figure 3c-e and Figure 4b). Another cytosolic Cu(I)-binding protein with a well-established role (18) in Cu homeostasis (delivering Cu(I) to *Bs*CopA) and a similar Cu(I) affinity (31) to *Bs*Csp3 (13) is *Bs*CopZ (see Figure 1a). As this was another potential source of Cu(I) for *Bs*CotA we tested to see if *Bs*CopZ could transfer Cu(I) to *Bs*CotA (Figure 3f and Figure 4a). After the incubation of apo-*Bs*CotA with Cu(I)-*Bs*CopZ for 48 h very little activity is observed, and Cu(I) transfer does not occur.

## Discussion

Herein we demonstrate that *Bs*Csp3 stores Cu(I) in the cytosol during sporulation for the Cu-requiring enzyme *Bs*CotA. This is not the only mechanism available to load *Bs*CotA with Cu as some activity is still observed in Δ*csp3 B. subtilis*. A possibility we considered was that the cytosolic Cu metallochaperone *Bs*CopZ, as well as transferring Cu(I) to *Bs*CopA, may also store cuprous ions in the cytosol, which could be transferred to *Bs*CotA (the Cu(I) affinity of *Bs*CopZ is similar to that of *Bs*Csp3 (13, 31)). This is supported by the suggestion that *Bs*CopZ could play a role in Cu(I) sequestration and recycling in *B. subtilis* (18) as less Cu accumulates in the Δ*copZ* strain compared to WT. Furthermore, the activity of CotA from a different *Bacillus* strain is enhanced when co-expressed in the cytosol of *E. coli* with its native CopZ (32). However, the *in vitro* studies reported here show that *Bs*CopZ cannot transfer Cu(I) to *Bs*CotA (Figure 3f and Figure 4a). The source(s) of Cu(I) for activating *Bs*CotA in the absence of *Bs*Csp3, and also at lower intracellular concentrations of the metal ion, remain(s) to be established. Regardless, the lack of Cu(I) transfer from *Bs*CopZ highlights the specificity of the *Bs*Csp3-*Bs*CotA interaction. This is essential in a cell as it ensures protein-mediated Cu(I) transfer to the correct destination, as observed for other Cu-homeostasis proteins (1, 33-39).

Considering the high Cu(I) affinity (13) of *Bs*Csp3 ((1.5 ± 0.4) × 10^17^ M^-1^), transfer of cuprous ions to *Bs*CotA has to occur via an associative mechanism (unassisted Cu(I) off-rates for *Bs*Csp3 can be estimated (14) to be ∼10^−9^ s^-1^). For the acquisition of such tightly bound Cu(I) to be possible, metalation must take place once the target enzyme has at least partially folded so the sites where Cu binds have formed. The T1 Cu site is closest to the surface of *Bs*CotA (the His497 ligand is solvent exposed), and is ∼12.5-15.5 Å from the Cu_3_ cluster (Figure 1c). Therefore, *Bs*Csp3 association at more than one location may be required to metalate all of the sites in folded *Bs*CotA. Published Cu(I) affinities of T1 Cu sites (40, 41) are (2.1-4.0) × 10^17^ M^-1^, similar to the average Cu(I) affinity of *Bs*Csp3 (13). Therefore, Cu(I) transfer from the storage protein to the enzyme should not be hindered thermodynamically (37, 40). To facilitate access to the more buried Cu_3_ cluster the protein may need to be partially unfolded. The MCO CueO from *E. coli* undergoes a transition from an ‘open’ non-metallated folded form with accessible Cu sites, to a more ‘closed’ conformation after Cu has bound (42). A similar change may occur in *Bs*CotA to facilitate Cu(I) loading by *Bs*Csp3.

The expression of both *Bs*Csp3 and *Bs*CotA are regulated by sigma factor K (SigK or σ^K^), which is produced after the forespore has been engulfed by the mother cell (Figure 5). Upregulation of the *csp3* gene occurs prior to *cotA* (Figure 1f), thus allowing the storage protein to acquire Cu(I) before production of the enzyme requiring the metal. As *Bs*CotA is one of the last proteins to be added to the spore coat (28), Cu(I) transfer from *Bs*Csp3 could occur during the very late stages of sporulation. Where *Bs*Csp3 acquires Cu(I) from is not known, but possible sources are Cu-enzymes that are not required by the spore, with the only currently known possibilities being COXs. The finding that *Bs*Csp3 provides Cu(I) to an enzyme requiring this metal ion suggests a similar role is performed by Csp3s in the cytosol of other bacteria. The novelty of this finding is further highlighted by there being only one currently known example of Cu acquisition by an enzyme from a partner protein in the cytosol. This is from the eukaryotic Cu metallochaperone CCS to the Cu,Zn-superoxide dismutase, which has been studied in considerable detail (1, 34, 35, 37-39, 43, 44).

**Figure 5.**
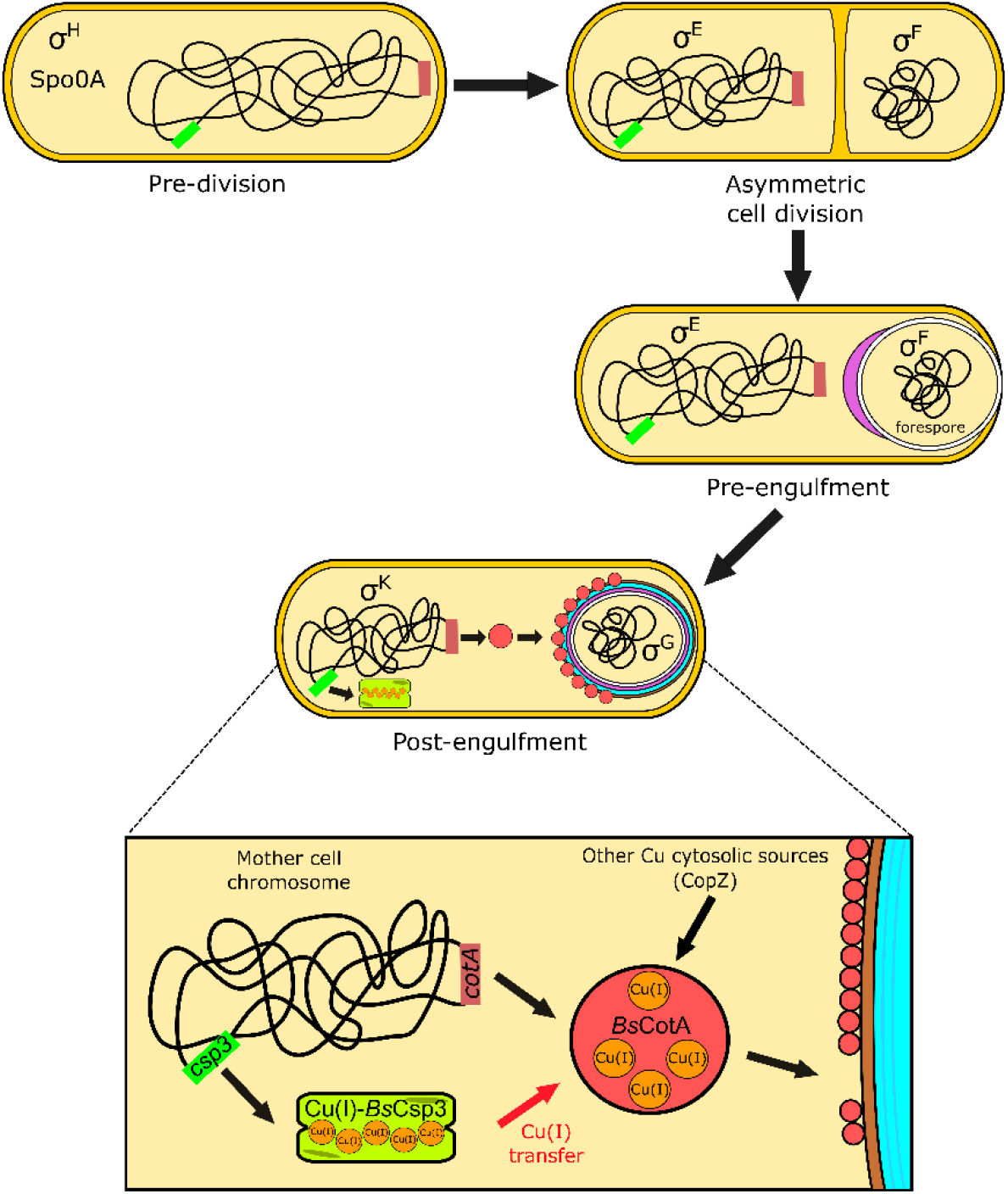
The proposed role of *Bs*Csp3 in Cu(I) acquisition by *Bs*CotA during sporulation in *B. subtilis*. The transcription factor Spo0A, along with σ^H^, initiates sporulation. A septum asymmetrically divides the cell into the forespore and mother cell, with σ^E^ and σ^F^, respectively, activated within these. The mother cell begins engulfment of the forespore and σ^E^ directs gene expression and initiation of spore coat (purple) formation. The expression of *Bs*Csp3 and *Bs*CotA now begins, promoted by σ^K^ (see Figure 1f) and coat assembly continues. We propose that *Bs*Csp3 acquires Cu(I) during this stage of sporulation, which is transferred to *Bs*CotA prior to insertion of the Cu-enzyme into the spore coat.

Added importance to understanding correct metalation of *Bs*CotA is provided by the observation that melanin formation interferes with the phagocytosis of pathogenic yeast, and is required to allow survival in macrophages (45). The related pigment produced by Cu-loaded *Bs*CotA is important for spore survival (25, 28), and this may include within a host. *Bacillus* spores, and particularly those from *B. cereus*, cause food poisoning and are a common contaminant in a range of foods (46, 47). The development of more effective inactivation approaches requires a better understanding of enzymes that help protect spores such as CotA. This includes establishing how they acquire essential cofactors including Cu ions.

## Materials and Methods

### Growth curves for WT and Δ*csp3 B. subtilis* at increasing Cu concentrations

WT and Δ*csp3 B. subtilis* 168 strains were obtained from the Bacillus Genetic Stock Centre library. The disrupted *csp3* gene (*yhjQ*) was amplified by PCR using genomic DNA from the Δ*csp3* strain with primers that hybridise ∼300 bp upstream and downstream of this region (Supplementary Table 3). The resulting fragment was sequenced with primers designed to hybridise ∼ 20 bp from the ends of the PCR product (Supplementary Table 3) and matches that of the erythromycin resistance gene. To test the influence of Cu on WT and Δ*csp3* strains, cultures were grown (agitation at 250 rpm) in LB media at 37 ºC overnight, diluted (∼100-fold) in LB and LB plus Cu(NO_3_)_2_ (0.5 to 2.0 mM). The absorbance at 600 nm was measured at regular intervals for up to 12 h, and also after 24 h growth. The Cu concentration in the stock solution used for these studies was regularly determined by atomic absorption spectrometry (AAS), as described previously (15).

### The construction of *B. subtilis* strains

To re-insert the *csp3* gene plus its ribosome binding site (RBS) into the Δ*csp3* strain, a region including an additional 28 bp at the 5’ end was amplified from *B. subtilis* 168 genomic DNA by PCR using primers; rbs_BsCsp3-F and rbs_BsCsp3-R (Supplementary Table 3). The product was cloned into pGEM-T (Promega) and the resulting *rbs_csp3* fragment sub-cloned into pDR111, which possesses the IPTG-inducible *P*_*hyerspank*_ promoter (48), using HindIII and NheI to generate pDR111_rbs_csp3. To obtain a strain possessing an IPTG-inducible copy of the *csp3* gene (complemented Δ*csp3*), Δ*csp3 B. subtilis* was transformed with pDR111_rbs_csp3. Selection was achieved using spectinomycin (50 μg/mL) and successful integration into the chromosomal *amyE* (α-amylase) gene identified by growing on LB agar containing 1% starch and staining with iodine (49). A strain of Δ*csp3 B. subtilis* in which the region of pDR111 lacking the *csp3* gene was integrated into the genome (control Δ*csp3*) was also generated. The size of the fragment incorporated was confirmed by PCR using the primers pDR111_int_F and pDR111_int_R (Supplementary Table 3).

### The production of *B. subtilis* spores

WT, Δ*csp3*, complemented Δ*csp3* and control Δ*csp3* strains were grown overnight (agitation at 250 rpm) in 20 mL Difco sporulation media (DSM). Cultures were diluted 50-fold into 200 mL DSM in a single 1 L Erlenmeyer flask and grown until the absorbance at 580 nm reached ∼0.5. This was split into four 50 mL cultures, each in a 250 mL Erlenmeyer flask, and 50 μM Cu(NO_3_)_2_ and 1 mM IPTG added when required. The cultures were grown (agitation at 250 rpm) at 37 °C for 48 h and absorbance values at 580 nm measured at regular intervals. To purify spores (50) cultures were centrifuged (4 °C) for 10 min at 5,000 g, pellets re-suspended in 50 mM Tris pH 7.2 plus 50 μg/mL lysozyme and incubated at 37 °C for 1 h. After incubation and further centrifugation (4 °C) for 10 min at 5,000 g, pellets were washed once in sterile MilliQ water and centrifuged. The pellets were re-suspended in 0.05% SDS by vortexing, centrifuged (4 °C) for 10 min at 5,000 g and subsequently washed three times with sterile MilliQ water. The purified spore stocks were verified by PCR (for example, Supplementary Figure 7) using primers listed in Supplementary Table 3 and stored at 4 °C.

### *Bs*CotA activity of purified spores

For kinetic measurements of *Bs*CotA activity, purified spores from the WT, Δ*csp3*, complemented Δ*csp3*, and control Δ*csp3* strains were diluted with MilliQ water to give an absorbance at 580 nm of ∼1.2 (measured accurately), and heated at 65 °C for 1 h prior to use. To determine the colony forming units per mL (CFUs/mL) for this suspension a 5 × 10^5^-fold dilution in LB was plated (100 μL) onto LB agar. The plates were incubated at 37 °C overnight and colonies counted. An aliquot of the heat-treated spore suspension (100 μL) was added to 900 μL of 100 mM citrate-phosphate buffer pH 4.0 plus 0.1-2.4 mM ABTS, and the absorbance at 420 nm (ε = 3.5 × 10^4^ M^-1^cm^-1^) measured for 5 min at 37 °C (Supplementary Figure 3). A control using 100 μL of buffer was also measured and showed no change in absorbance at 420 nm. The initial velocity (*V*_0_; typically reported in units of μM/min/CFUs/mL) was calculated, and plots of *V*_0_ against ABTS concentration (Figure 2 and Supplementary Figure 5) were fit to the Michaelis-Menten equation to determine *V*_max_ (the maximum rate) and *K*_M_ (the Michaelis constant). Comparing *V*_max_ values calculated based on the absorbance at 580 nm of the heat-treated spore suspension, rather than using CFUs/mL, has no significant influence on the outcome of the study, but generally produces data with larger errors. The reactivity of heat-treated purified spores from WT and Δ*csp3 B. subtilis* with 2.4 mM ABTS was compared to that of the same spore suspension incubated with 250 μM Cu(NO_3_)_2_ for 30 min at room temperature.

### Cloning and purification of *Bs*CotA

The *cotA* gene was amplified from *B. subtilis* genomic DNA using primers CotA_1F and CotA_1R listed in Supplementary Table 3, and cloned into pGEM-T. After removing the NdeI site in the gene by QuickChange site-directed mutagenesis (with primers CotA_2F and CotA_2R, Supplementary Table 3), the product was excised with NdeI and BamHI and re-cloned into pET11a. *Bs*CotA was overexpressed in BL21 *E. coli* (100 μM IPTG) grown at 20 °C for 24 h. The protein was purified using a modified version of a published procedure (26). Cells from 0.5 to 1.0 L of culture were resuspended in 20 mM Tris pH 8.5, sonicated and centrifuged at 40,000 g for 30 min. The supernatant was diluted five-fold in 20 mM Tris pH 8.5 and loaded onto a HiTrap Q HP column (1 mL) equilibrated in the same buffer. Proteins were eluted with a linear NaCl gradient (0-500 mM, total volume 50 mL) and fractions analysed using 18% SDS-PAGE. *Bs*CotA-containing fractions were diluted with 20 mM Tris pH 7.6 and loaded onto a HiTrap SP HP column (5 mL) and eluted with a linear NaCl gradient (0-500 mM, total volume, 200 mL). The *Bs*CotA-containing fractions were heated at 70 °C for 30 min (*Bs*CotA is a highly thermostable enzyme (26)), centrifuged at 40,000 g for 30 min, and the supernatant exchanged into 20 mM HEPES pH 7.5 plus 200 mM NaCl for further purification on a Superdex 75 10/300 GL gel-filtration column. Purified *Bs*CotA had very little Cu or Zn(II) (the latter, a metal ion that commonly binds to Cu proteins when overexpressed in the cytosol of *E. coli*) associated with it when analysed by AAS (12, 13), and showed almost no ABTS oxidation activity. Samples (∼3-12 μM, quantified using an ε value of 84,739 M^-1^cm^-1^ at 280 nm (51)) were incubated with 250 μM Cu(NO_3_)_2_, thoroughly exchanged and subsequently washed with a low concentration (∼10 μM) of ethylenediaminetetraacetic acid (EDTA). This gave rise to Cu-loaded *Bs*CotA with a *k*_cat_ value of 20-40 s^-1^ for the oxidation of ABTS, compared to a literature value of 22 s^-1^ for Cu-enzyme produced using a similar procedure (51).

### Purification of *Bs*CopZ and sample preparation

*Bs*CopZ is purified with a small amount of Zn(II) bound, as described previously (13). Samples were therefore incubated with >10 equivalents of EDTA for 1 h and exchanged with 20 mM HEPES pH 7.5 plus 200 mM NaCl. The resulting protein had no Zn(II) associated with it and was reduced with dithiothreitol under anaerobic conditions and desalted as described previously (12, 13).

### Analysing Cu(I) transfer from recombinant *Bs*Csp3 and *Bs*CopZ to *Bs*CotA

*Bs*Csp3 binding ∼18 equivalents of Cu(I) was prepared by adding the appropriate amount of a buffered solution of Cu(I) in an anaerobic chamber (Belle Technology, O_2_ << 2 ppm) to apo-protein in 20 mM HEPES pH 7.5 plus 200 mM NaCl, quantified using the 5,5’-dithiobis(2-nitrobenzoic acid) (DTNB) assay (12, 13). Fully-reduced *Bs*CopZ was also quantified using the DTNB assay and loaded with ∼0.8 equivalents of Cu(I) under anaerobic conditions. Cu(I)-*Bs*Csp3 (∼3 μM binding ∼53-54 μM Cu(I)) was mixed with ∼1.3 μM of as-isolated *Bs*CotA, requiring ∼5.6 μM Cu(I) to occupy all Cu sites. Cu(I)-*Bs*CopZ (∼50-53 μM binding ∼40-42 μM Cu(I)) was separately added to ∼1 μM of as-isolated *Bs*CotA, requiring ∼4 μM Cu(I) to fill all Cu sites. Mixtures were incubated at room temperature in the anaerobic chamber for up to 48 h. A similar concentration of *Bs*CotA was also incubated anaerobically with Cu(I), prepared as described above, a Cu(NO_3_)_2_ solution (final concentrations of ∼56-58 μM), buffer (20 mM HEPES pH 7.5 plus 200 mM NaCl) alone, and also with Cu(NO_3_)_2_ in air. To measure activity, 10 μL of each mixture was added to 990 μL of aerated 100 mM citrate-phosphate buffer pH 4.0 plus 2.4 mM ABTS, and the absorbance at 420 nm measured for up to 5 min at 37 °C (Figure 3). The removal of Cu(I) by BCS (∼2.5 mM) was analysed for Cu(I)-*Bs*Csp3 (∼0.8-1.2 μM plus ∼18 equivalents of Cu(I)) samples used for the transfer experiments, both in the absence (folded *Bs*Csp3) and presence (unfolding conditions) of guanidine-HCl (6.64 M) (12, 13). The absorbance increase at 483 nm due to formation of [Cu(BCS)_2_]^3-^(ε = 12,500 M^-1^cm^-1^) (31) was measured over time at 22 °C in 20 mM HEPES pH 7.5 plus 200 mM NaCl (Supplementary Figure 6).

## Acknowledgments

We thank BBSRC for funding (grant BB/K008439/1 to C.D.). We are grateful to Newcastle University for an Overseas Research Scholarship (ORS) award to J.L.. We also thank Prof. Heath Murray for help with generating *B. subtilis* mutant strains, Dr. Mark Harrison for cloning the *cotA* gene and Dr. Gianpiero Landolfi for purifying *Bs*CopZ.

## Author contributions

C.D. and J.L conceived the project and designed the experiments. J.L. performed the experiments and analysed data with help from C.D. C.D. wrote the manuscript with help from J.L.

## Competing interests

The authors declare no competing interests.

## Additional information

Supplementary information is available for this paper.

## SUPPLEMENTARY INFORMATION

**Supplementary Figure 1.**
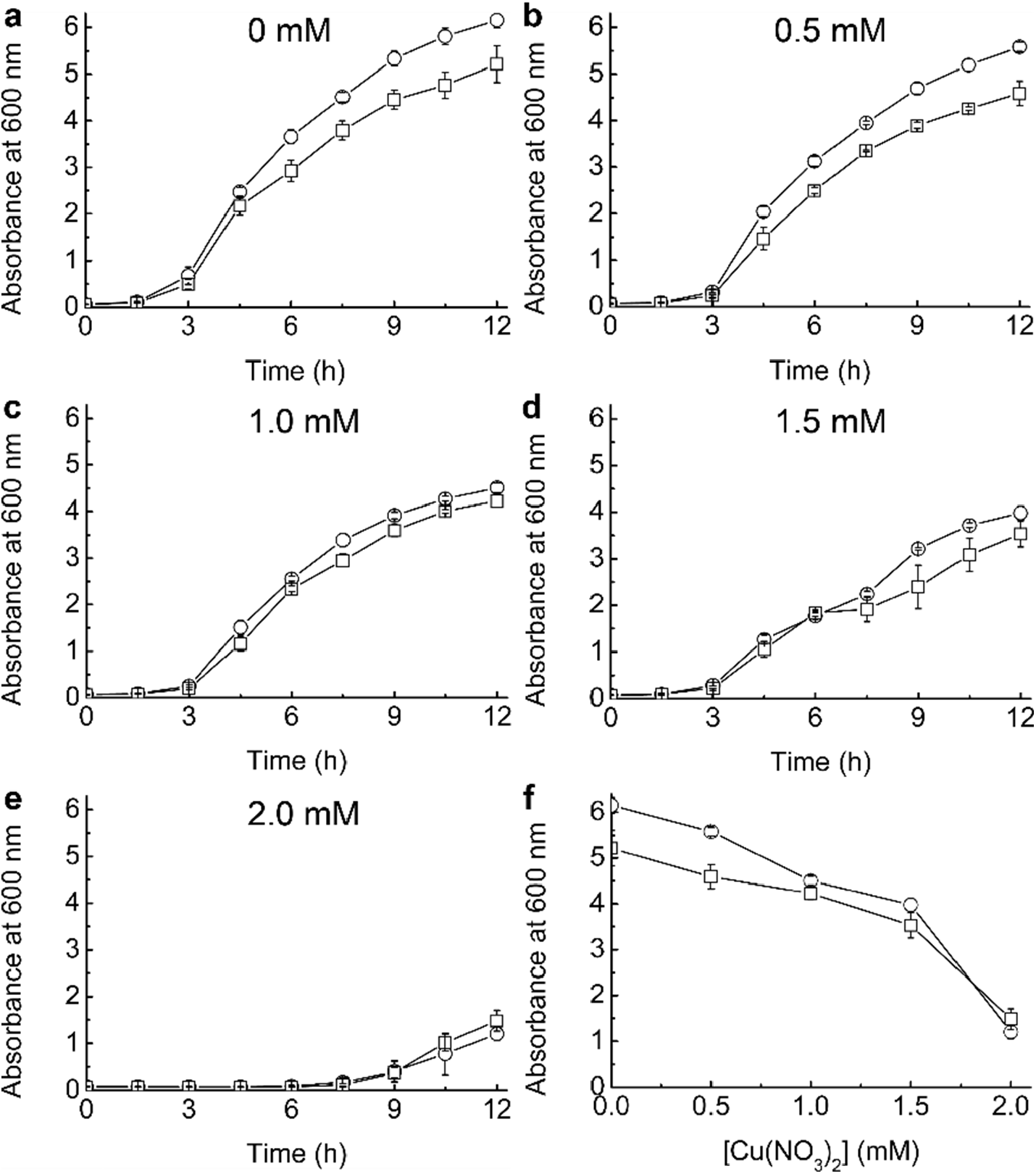
The influence of Cu on the growth of WT and Δ*csp3 B. subtilis*. Growth (37 °C) of WT (circles) and Δ*csp3* (squares) *B. subtilis* in LB media plus 0 (a), 0.5 (b), 1.0 (c), 1.5 (d), and 2.0 (e) mM added Cu(NO_3_)_2_. The data obtained at 12 h is compared in (f), and in all cases averages and standard deviations from three independent growth experiments are shown. These results are similar to those we reported previously (13), albeit the influence on growth starts to be observed at a Cu(II) concentration in the media that is ∼0.5 mM lower in this work (we have quantified Cu levels in the stock solutions used herein by AAS, but did not previously (13)). The absorbance of these cultures were also measured at 24 h, and at up to 1.0 mM added Cu(NO_3_)_2_ a significant decrease was observed compared to the value at 12 h for both strains (consistent with previous data (13)). At higher added Cu(NO_3_)_2_ concentrations, the growth data showed no consistent pattern beyond 12 h. This differs to the enhanced cell death that was seen previously for Δ*csp3 B. subtilis* after 12 h growth at reported media Cu concentrations of 1.5-2.0 mM (13).

**Supplementary Figure 2.**
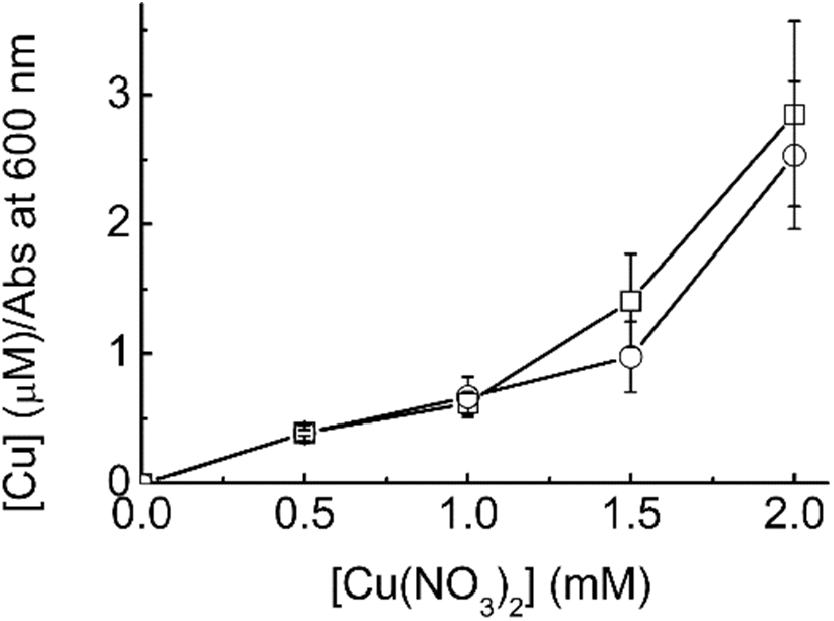
The influence of Cu levels in media on Cu accumulation by WT and Δ*csp3 B. subtilis*. The intracellular Cu concentrations for WT (circles) and Δ*csp3* (squares) *B. subtilis* grown for 12 h in LB media plus increasing amounts of added Cu(NO_3_)_2_. The data shown (average values and standard deviations) were measured for only two of the independent growth experiments, but the trend is clear.

**Supplementary Figure 3.**
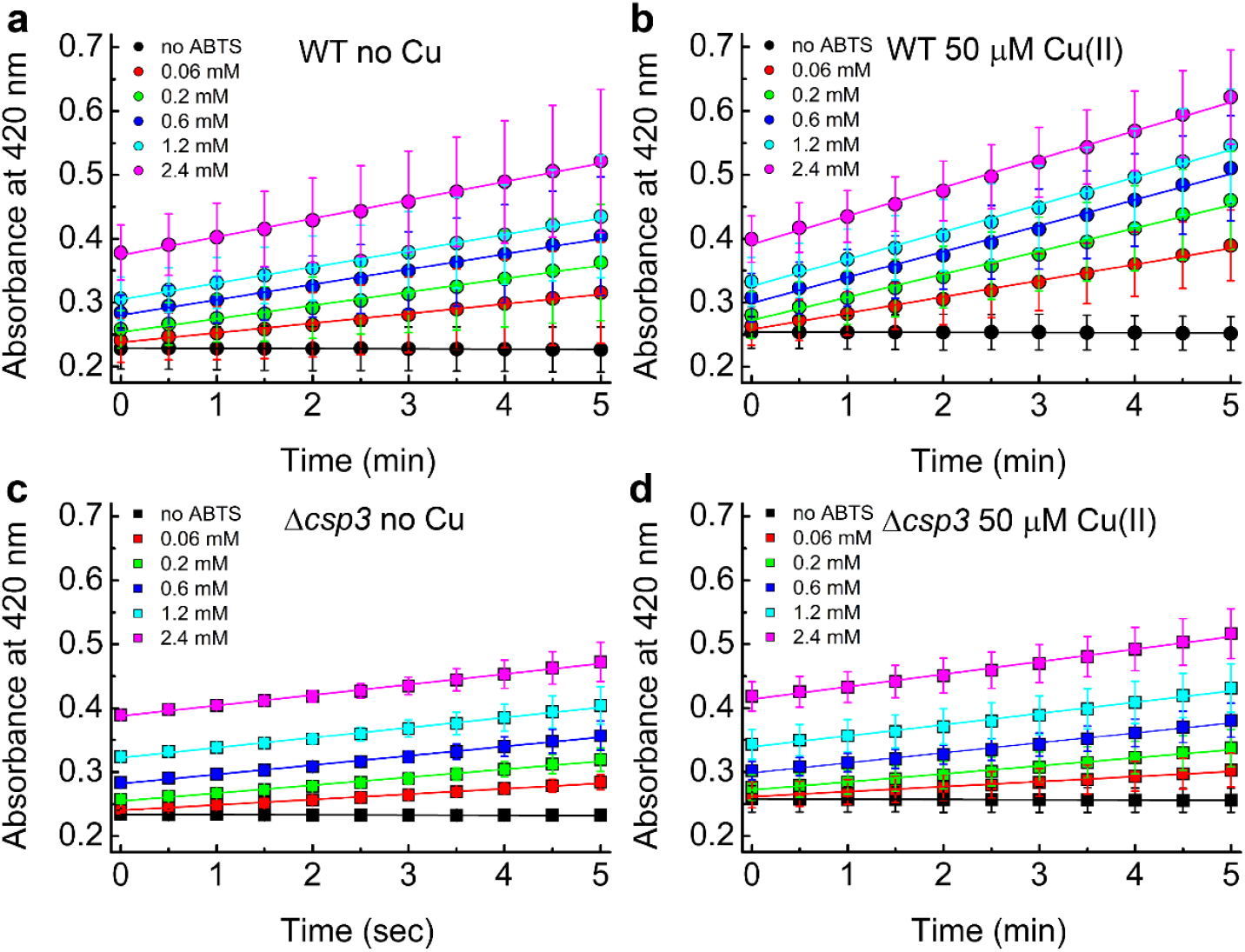
The influence of Cu levels and deleting the *csp3* gene on *Bs*CotA activity in *B. subtilis* spores. Plots of absorbance at 420 nm against time at different concentrations of ABTS (indicated) for spores from WT (a) and (b) and Δ*csp3* (c) and (d) *B. subtilis*. The data in (a) and (c) are from spores obtained in DSM without added Cu, whilst 50 μM Cu(NO_3_)_2_ was added for (b) and (d). The reactions with ABTS were measured in 100 mM citrate-phosphate buffer pH 4.0 and the initial rates (averages from three different sets of spores with error bars showing standard deviations) we used for Figure 2a (WT) and 2b (Δ*csp3*).

**Supplementary Figure 4.**
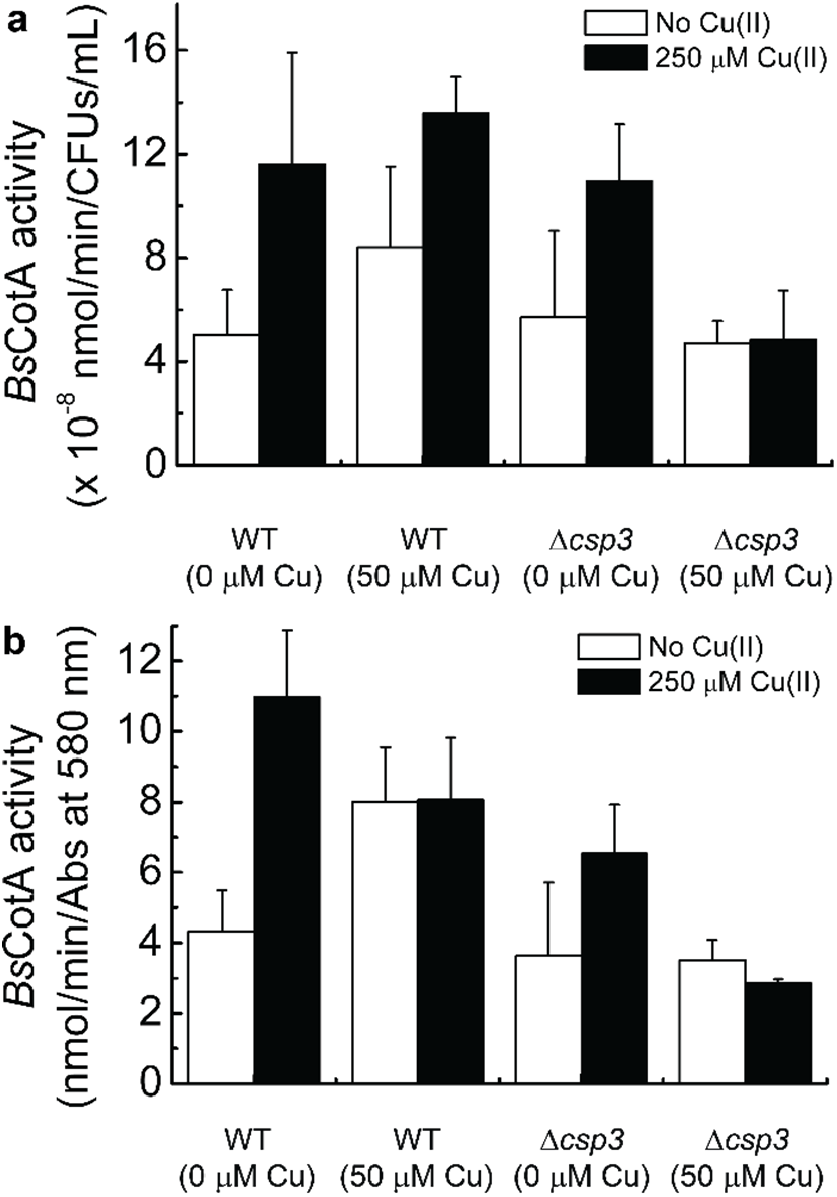
Testing purified spores for enhanced *Bs*CotA activity after incubation with Cu(II). The influence of incubating spores in 250 μM Cu(NO_3_)_2_ from WT and Δ*csp3 B. subtilis* grown either in the absence or presence (50 μM) of Cu(NO_3_)_2_ on their *Bs*CotA activity. The activity units are reported based on both CFU values (a) and the absorbance (Abs) at 580 nm (b) to quantify spores. Average values and standard deviations from three independent experiments on the same spores used for the data in Figure 2a,b are shown, and the values are listed in Supplementary Table 1. The reactions with ABTS were measured in 100 mM citrate-phosphate buffer pH 4.0.

**Supplementary Figure 5.**
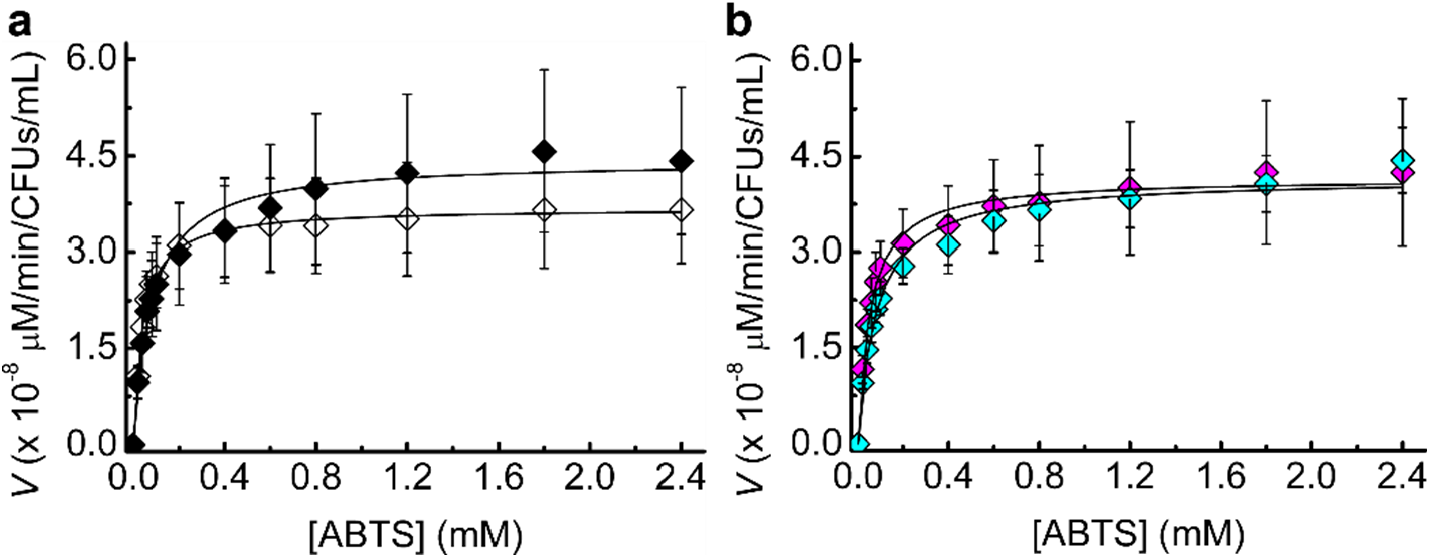
The influence of Cu levels and IPTG on *Bs*CotA activity in the control Δ*csp3 B. subtilis* strain. Michaelis-Menten plots of *Bs*CotA activity for heated purified spores from the control Δ*csp3* strain. (a) The analysis of spores produced in media plus 0 (open symbols) and 50 (black filled symbols) μM added Cu(NO_3_)_2_ with no added IPTG. (b) Data plus 1 mM IPTG and 0 (magenta) and 50 (cyan) μM Cu(NO_3_)_2_. Averages and standard deviations from kinetic measurements in 100 mM citrate-phosphate buffer pH 4.0 using three different sets of spores are shown.

**Supplementary Figure 6.**
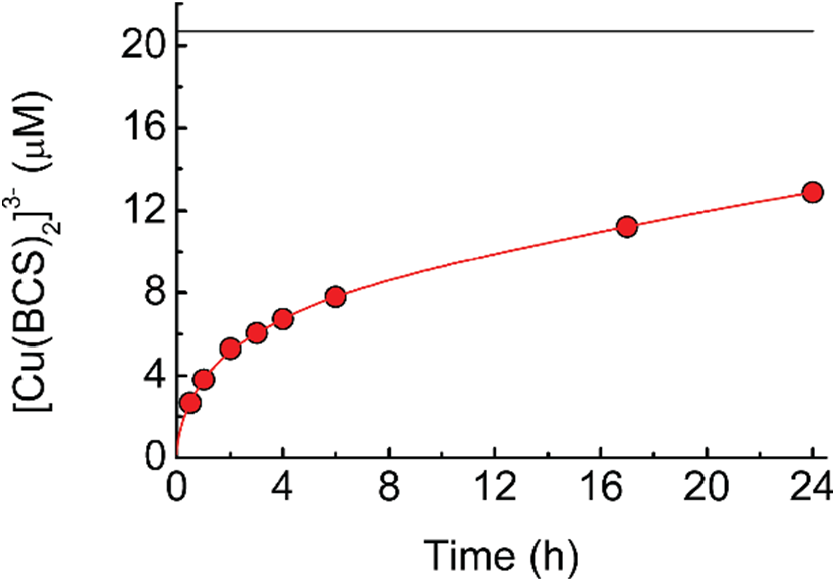
The removal of Cu(I) from *Bs*Csp3 by the high-affinity ligand BCS. A plot of [Cu(BCS)_2_]3-concentration against time for *Bs*Csp3 (1.08 μM) plus 18.0 equivalents of Cu(I) mixed with 2.5 mM BCS in 20 mM HEPES pH 7.5 plus 200 mM NaCl (red line). The data at 0.5, 1, 2, 3, 4, 6, 17 and 24 h, which correspond to times at which *Bs*CotA activity was measured (Figure 4), are indicated by red circles. Average percentage removal compared to that for the unfolded samples, and the standard deviations, from three independent experiments are shown in Supplementary Table 2. The black line indicates the same experiment but with 6.64 M guanidine-HCl present in the buffer. This helps unfold the protein and removal is much faster giving the end point for the reaction (the value shown was obtained after incubation for 2 h).

**Supplementary Figure 7.**
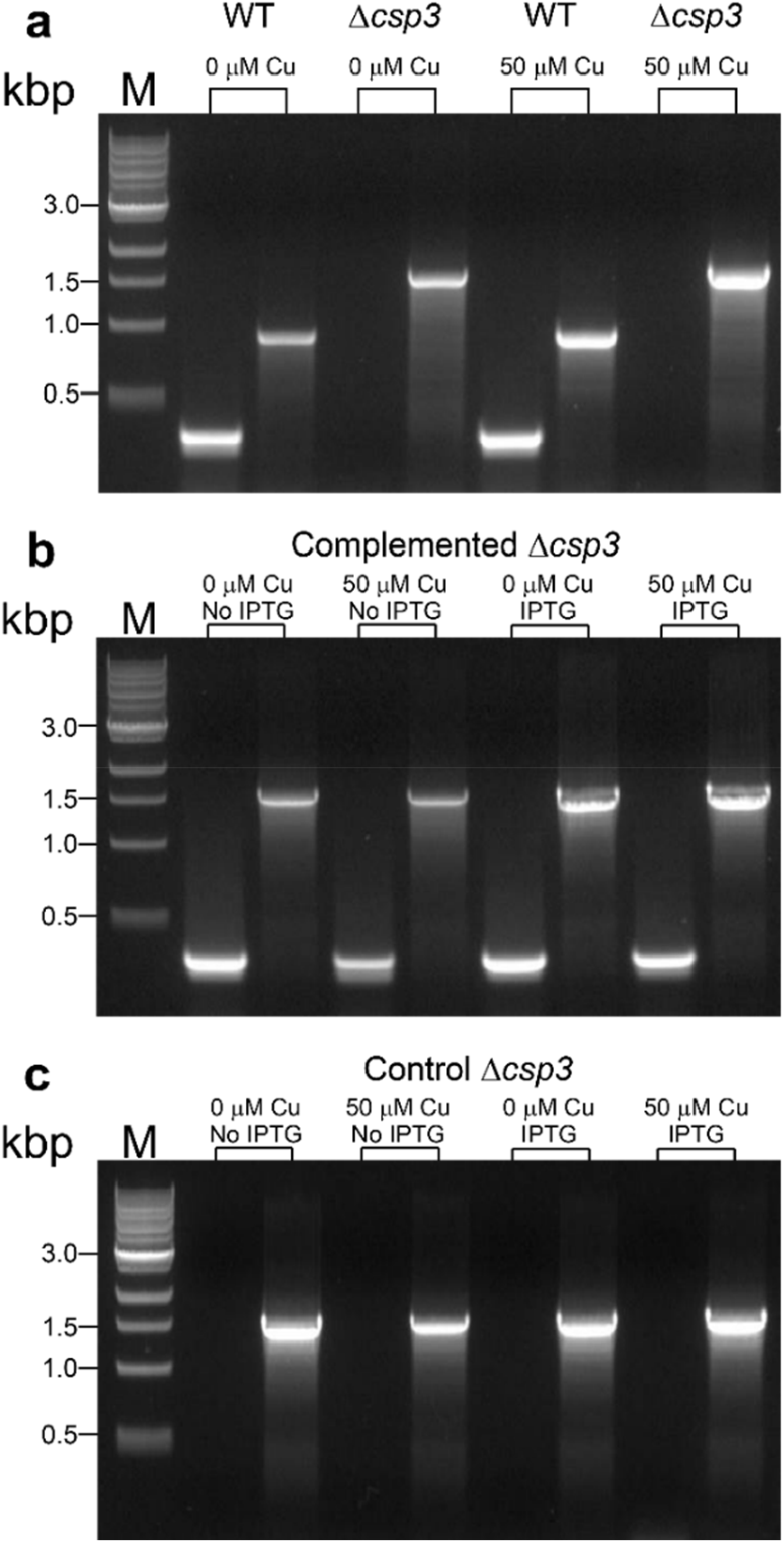
PCR analyses of purified spore stocks from the *B. subtilis* strains used in this study. (a) The rbs_BsCsp3-F and rbs_BsCsp3-R primers (Supplementary Table 3) give a product of 373 bp for the WT strain, whilst BsCsp3+300-F and BsCsp3+300-R give fragments of 967 and 1663 bp for WT and Δ*csp3*, respectively. (b) Re-introduction of the *csp3* gene is confirmed in the complemented strain using the rbs_BsCsp3-F and rbs_BsCsp3-F primers (as well as testing with BsCsp3+300-F and BsCsp3+300-R), and the same primers are used to analyse the control Δ*csp3* strain (c). Analysis of spores used for the three independent experiments on all four strains shown in Figure 2 and Supplementary Figure 5 gave the same results.

**Supplementary Table 1.**
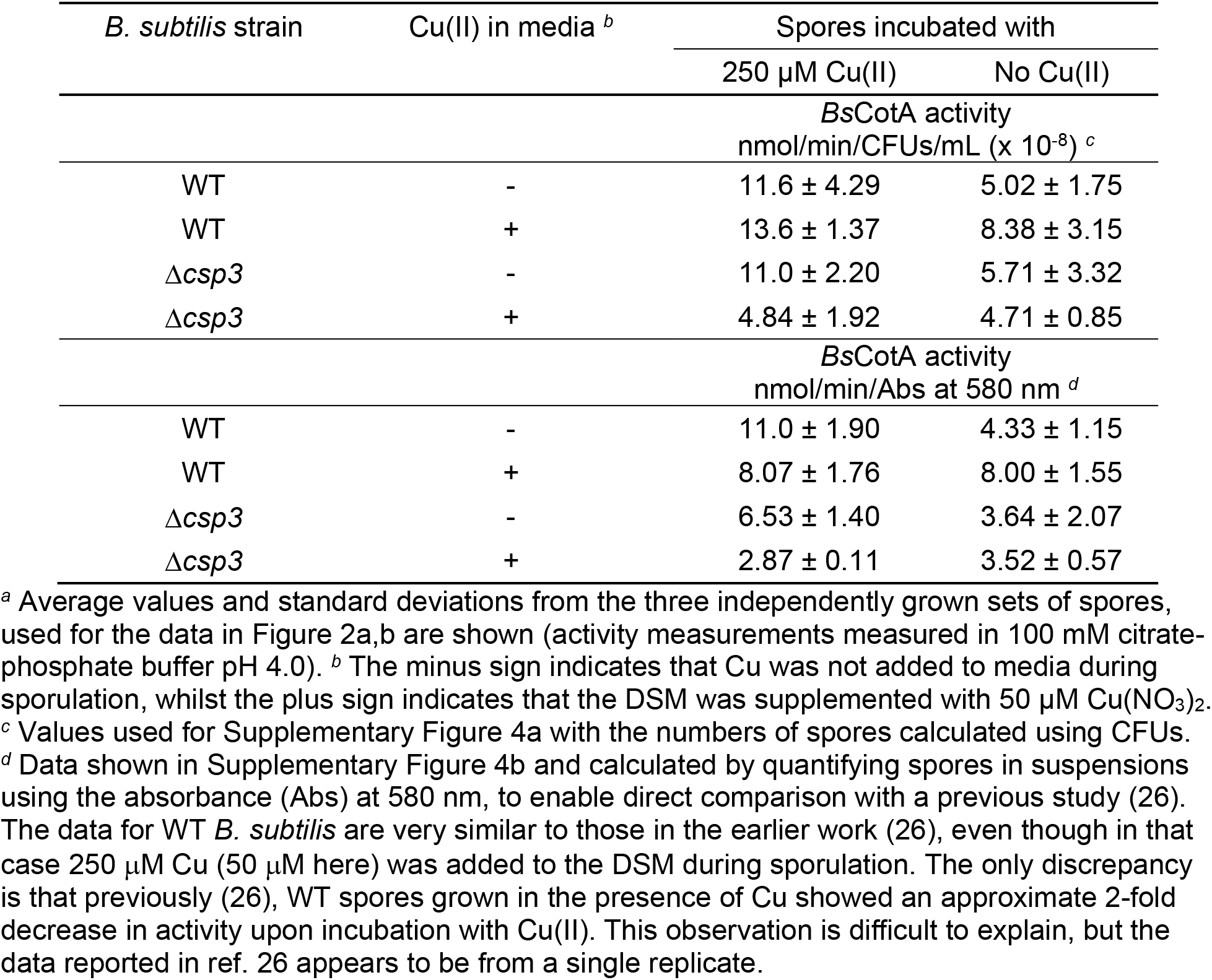
The influence of Cu concentration on the *Bs*CotA activity of purified spores from WT and Δ*csp3 B. subtilis a*

**Supplementary Table 2.** The removal of Cu(I) from *Bs*Csp3 by BCS over time *a,b*

| Time (h) | % Cu(I) removal <sup>c</sup> |
| --- | --- |
| 0.5 | 12.9 ± 0.54 |
| 1 | 18.0 ± 0.71 |
| 2 | 24.7 ± 1.33 |
| 3 | 28.3 ± 1.23 |
| 4 | 31.5 ± 1.39 |
| 6 | 36.6 ± 1.50 |
| 17 | 52.9 ± 2.31 |
| 24 | 60.7 ± 2.35 <sup>d</sup> |

<sup>a</sup> Shown are the average percentage removal of Cu(I) from *BsCsp3* by BCS over time (and standard deviations) from three independent experiments. <sup>b</sup> The amount of Cu(I) bound to the protein was determined by mixing the Cu(I)-*BsCsp3* sample with 2.5 mM BCS in the presence of 6.64 M guanidine-HCl in 20 mM HEPES pH 7.5 plus 200 mM NaCl, which unfolds the protein giving the maximum possible [Cu(BCS)<sub>2</sub>]<sup>3-</sup> concentration (the value used is after incubation for 2 h). <sup>c</sup> Percentage Cu(I) removal is determined using [Cu(BCS)<sub>2</sub>]<sup>3-</sup> concentration/maximum [Cu(BCS)<sub>2</sub>]<sup>3-</sup> concentration × 100. <sup>d</sup> On one occasion the experiment was analysed up to 48 h with 70% Cu(I) removal observed.

**Supplementary Table 3.**
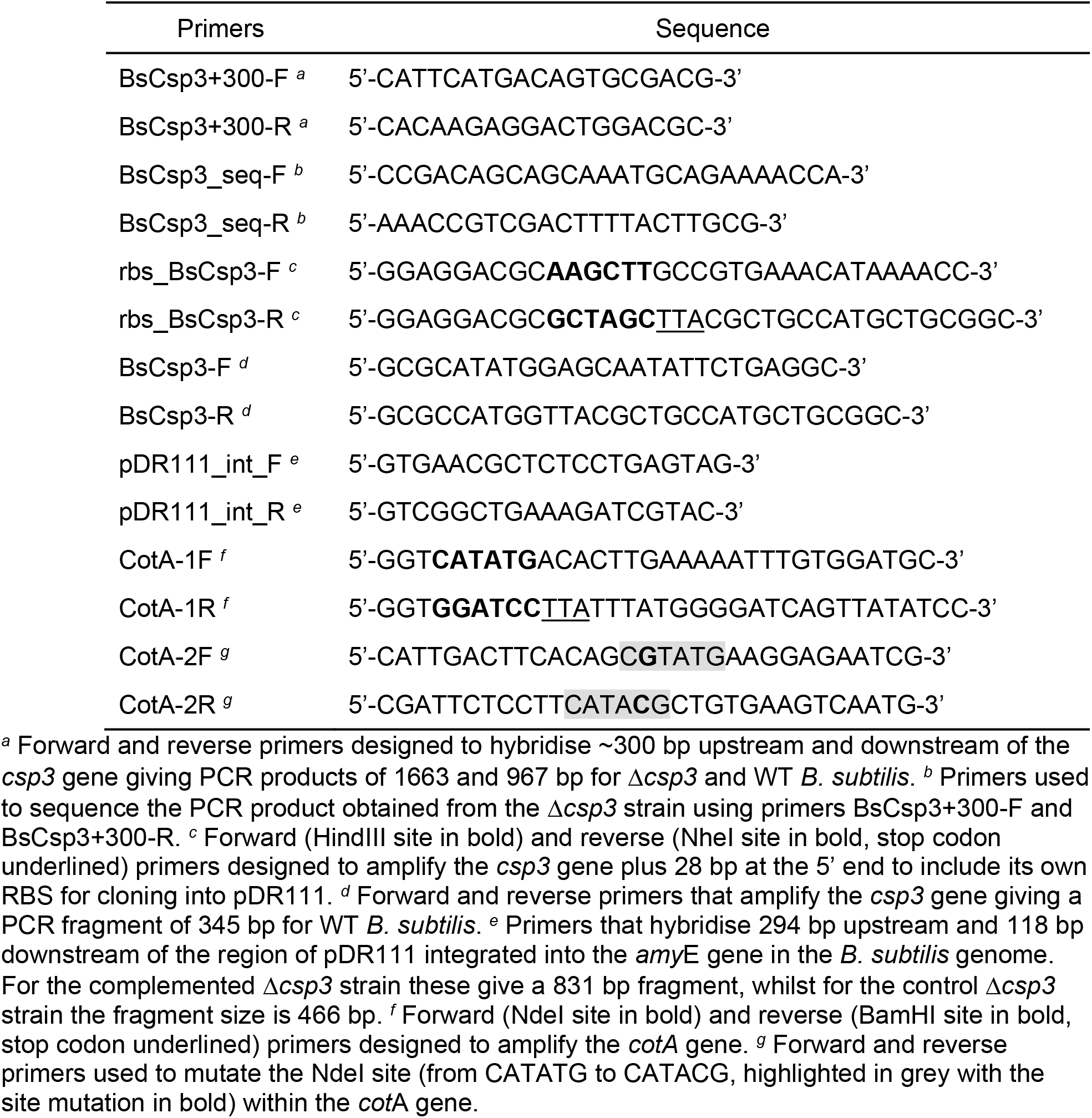
Primers used in this study

## Notes

### Competing Interest Statement

The authors have declared no competing interest.

